# High efficiency excitation energy transfer in biohybrid quantum dot–bacterial reaction center nanoconjugates

**DOI:** 10.1101/2021.04.20.440209

**Authors:** Giordano Amoruso, Juntai Liu, Daniel W. Polak, Kavita Tiwari, Michael R. Jones, Thomas A. A. Oliver

## Abstract

Reaction centers (RCs) are the pivotal component of natural photosystems, converting solar energy into the potential difference between separated electrons and holes that is used to power much of biology. RCs from anoxygenic purple photosynthetic bacteria such as *Rhodobacter sphaeroides* only weakly absorb much of the visible region of the solar spectrum which limits their overall light-harvesting capacity. For *in vitro* applications such as bio-hybrid photodevices this deficiency can be addressed by effectively coupling RCs with synthetic lightharvesting materials. Here, we studied the time scale and efficiency of Förster resonance energy transfer (FRET) in a nanoconjugate assembled from a synthetic quantum dot (QD) antenna and a tailored RC engineered to be fluorescent. Time-correlated single photon counting spectroscopy of biohybrid conjugates enabled the direct determination of FRET from QDs to attached RCs on a time scale of 26.6 ± 0.1 ns and with a high efficiency of 0.75 ± 0.01.

---

Purple bacteria such as *Rhodobacter (Rba.) sphaeroides* are among the simplest organisms capable of photosynthesis. These bacteria inhabit aquatic environments where they use pigment-protein complexes to absorb spectrally-filtered sunlight to power growth and reproduction.^1,2^To avoid competing with chlorophyll-containing oxygenic phototrophs, bacteria such as *Rba. sphaeroides* capture primarily near-infrared and blue/near-ultraviolet solar wavelengths using bacteriochlorophyll and carotenoid pigments.^1,2^In reaction center-light harvesting 1 (RC–LH1) complexes, a central RC electron transfer (ET) protein is encircled by an LH1 light harvesting pigment-protein. These RC–LH1 complexes are in turn surrounded in the membrane by peripheral LH2 light harvesting complexes.^3–5^Through sequential ultrafast electronic energy transfer (EET) steps, the initially-formed excited electronic state of LH2 or LH1 pigments is funneled to a RC where the excitation energy is used to drive rapid and efficient charge separation. The quantum yield of EET is very high because energy migration to the RC is fast (tens of picoseconds) compared to the excited state lifetime of bacteriochlorophyll (a few nanoseconds).^1,2^RCs perform the crucial energy transduction and stabilization processes of photosynthesis with a quantum yield that is near to unity,^6^efficiently separating charge between a pair of bacteriochlorophylls on one side of the membrane that act as the primary electron donor (termed P), and a ubiquinone on the opposite side (see Figure 1(a)).^7–9^

**Figure 1.**
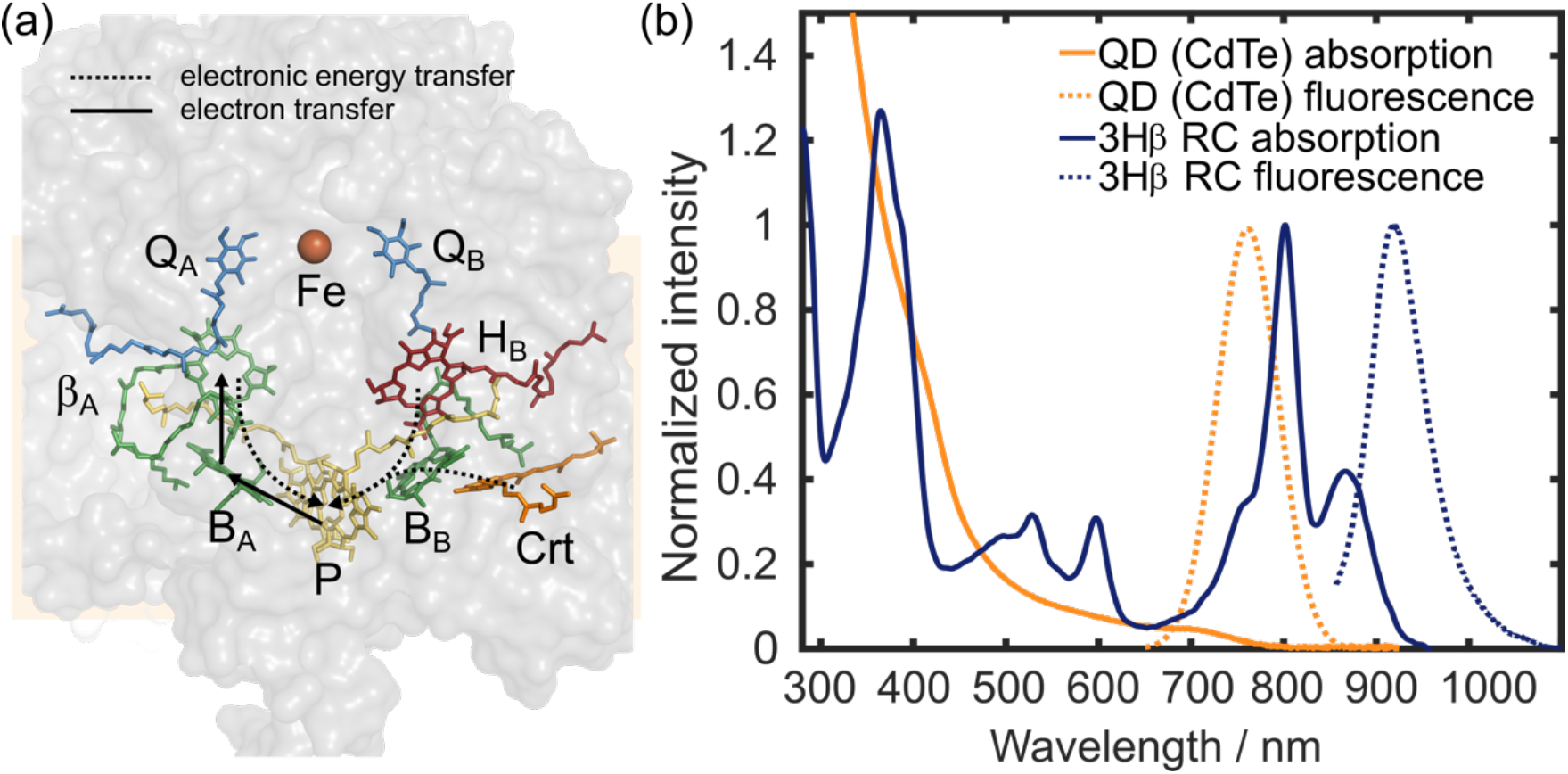
(a) Structural model of the 3Hβ RC protein (gray) highlighting the arrangement of light absorbing pigments (colored). One of the 3Hβ mutations substitutes the native A-branch bacteriopheophytin (H_A_) with a bacteriochlorophyll (labelled β_A_). Two different EET pathways are expected on the two sides of the RC (dashed arrows). For the B-branch (right) EET occurs between the bacteriopheophytin (H_B_) or the 15 *cis-cis’;* spheroidenone carotenoid (Crt) to a monomeric bacteriochlorophyll (B_B_), and thence to the special pair of bacteriochlorophylls (P). Alternatively, on the A-branch (left) the EET path is from bacteriochlorophyll β_A_ to bacteriochlorophyll B_A_, and then on to P. In the WT RC, formation of the excited singlet state of P (P*) triggers charge separation to sequentially form the radical pairs P^+^B_A_^−^, P^+^H_A_^−^and P^+^Q_A_^−^. It is anticipated in the 3Hβ RC only a very minor percent (< 1%) undergo charge separation to form a mixed P^+^(B_A_β_A_)^−^ radical pair. (b) Absorption and fluorescence spectra (normalized to the maximum intensity in 700–950 nm region) of the 3Hβ RC and QD components of the nanoconjugates.

These very high quantum efficiencies make RCs and RC–LH1 complexes attractive components for alternative solar energy conversion technologies employing environmentally-benign materials.^10^However, their limited capacity to harvest sunlight across much of the visible region of the available solar spectrum has obvious consequences for their performance in biohybrid photodevices, with relatively low external quantum efficiencies of photocurrent generation across green and red wavelengths in particular.^10–17^One way to address this drawback is to mimic the role of natural antenna pigment-proteins through conjugation of the RC with a non-native light harvesting system such as a synthetic quantum dot (QD),^18–24^a synthetic dye,^25–28^or a protein with complementary pigmentation.^29,30^This enables enhanced light collection over parts of the electromagnetic spectrum where the absorbance of the native RC pigmentation is weak.

Recently, Liu and co-workers characterized a biohybrid nanoconjugate in which *Rba. sphaeroides* RCs engineered with a poly-histidine tag were tethered to the surface of watersoluble CdTe QDs.^31,32^In these nanoconjugates the QDs served as both an assembly hub and an artificial light harvesting antenna for the RCs. On mixing, the His-tag bound the RCs to the surface of the QD with high affinity. Quenching of QD fluorescence upon increasing the ratio of tethered RCs per QD provided indirect proof of Förster resonance energy transfer (FRET) from the QDs to the attached RCs. As wild-type (WT) RCs are highly efficient at charge-separation, with a fluorescence quantum yield of virtually zero, a second set of nanoconjugates were constructed using a weakly fluorescent RC mutant (VL157R), that was unable to carry out charge separation due to the absence of one of the bacteriochlorophylls that make up the primary electron donor.^31^In this second set, the intensity of RC fluorescence peaking at 801 nm rose as the RC:QD ratio increased, mirroring a progressive decrease in overlapping QD fluorescence at 750 nm and providing further evidence for FRET between the QDs and the bound RCs.^31^A variety of other RC mutants were used in this prior study to explore the effect of changing the overlap integral between QD fluorescence and RC absorption on QD fluorescence quenching.

A key aspect of natural photosynthesis is the very high quantum efficiency of EET even in very extensive antenna systems. One potential way to establish the time scale and efficiency of EET in these synthetic biohybrid nanoconjugates is to use time-correlated single photon counting (TCSPC) spectroscopy.^33^This technique has previously been used to interrogate FRET dynamics between a natural plant light harvesting complex and QD hybrid.^34^However, in previous studies of WT RCs electrostatically bound to QDs it would not have been possible to apply TCSPC to the non-fluorescent WT RC energy acceptor.^18–20^Furthermore, although the VL157R RC mutant used in our previous work is fluorescent with a maximum at 801 nm,^31,35^TCSPC characterization of fluorescence changes at this wavelength caused by EET from associated QDs would be hampered by strongly overlapping fluorescence from the 750 nm emitting QD donors. The 801 nm fluorescence in VL157R RC is from the retained three monomeric bacteriochlorophylls, this RC having lost the red-shifted absorbance at ∼870 nm (and associated very weak fluorescence) from the pair of P bacteriochlorophylls normally present in the WT RC.^35^

To use TCSPC to determine the efficiency of EET in RC–QD nanoconjugates, a new RC mutant was engineered, 3Hβ (Figure 1(a)), that retains a normal long-wavelength primary electron donor but has a drastically-reduced quantum yield for charge separation (see SI, Section 1). This was achieved by combining a triple mutation LL131H + LM160H + FM197H,^36–38^with the so-called ‘ β-mutation’ LM214H.^39,40^The triple mutation adds three hydrogen bonds between the protein scaffold and the two bacteriochlorophylls that make up P, raising its mid-point potential for one electron oxidation by around 260 mV but without markedly affecting its absorption spectrum.^37^This has the effect of raising the free energy associated with the first radical pair formed during charge separation, P^+^B_A_^−^, above that of the excited singlet state of P (P*), thus severely slowing primary charge separation (causing a ∼50% decrease in quantum yield) and enhancing the fluorescence of P* through an extension of its lifetime from ∼3 ps to ∼50 ps.^38^The ‘ β-mutation’ replaces the bacteriopheophytin that is the second acceptor during charge separation (H_A_) with a bacteriochlorophyll (βA) (Figure 1a).^39,40^ In isolation this mutation renders the first (B_A_) and second (β_A_) electron acceptor isoenergetic, enabling formation of an admixture of the P^+^B_A_^−^ and P^+^ β_A_^−^ radical pairs. This slows the rate of charge separation by a factor of two and lowers its quantum yield to ∼60%. Formation of a mixed P^+^(B_A_β_A_)^−^ state both greatly reduces the probability of further electron transfer to the A branch quinone and strongly promotes the reverse electron transfer reaction to re-form P*.^39,40^ In the combined 3Hβ mutant it is expected that the quantum yield of charge-separated species should be drastically lower than for WT RCs, and strongly enhanced P* fluorescence should be observable with a maximum at ∼910 nm.^41^

Figure 1(b) shows the absorption and emission spectra of the QDs and His-tagged 3Hβ RCs used to assemble nanoconjugates in the current study (see SI, Section 2). As in previous work,^31,32^water-soluble CdTe QDs of 6.5 nm diameter were used as they have broad absorbance (solid orange line in Figure 1(b)) that spans the gap between the strong near-IR *Q*_y_ and near-UV Soret absorbance bands of the RC, and a fluorescence profile (dashed orange line) that overlaps well with RC absorption (solid dark blue line). The absorption spectrum of the 3Hβ RC was assigned by comparison to that of WT RCs (see overlay in Fig. S1), the main effect of the four mutations being a red-shift of the *Q*_y_ band at 760 nm due to substitution of the H_A_ bacteriopheophytin with the β_A_ bacteriochlorophyll, and a broadening on the blue edge of the 802 nm band attributed to the two native monomeric RC bacteriochlorophylls.^39^Fluorescence from the P bacteriochlorophyll pair peaked at 910 nm (blue dashed line in Figure 1(b)) and was spectrally shifted and resolved from QD fluorescence (orange dashed line).

To explore whether 3Hβ RCs were capable of charge separation, ultrafast transient absorption (TA) studies of purified RCs were undertaken using 59 fs, 595 nm pump pulses and two different supercontinuum probes in the visible or near-infrared (see SI Section 4 for full experimental details). Pulses at 595 nm excited the strongly overlapping *Q*_x_ bands of the five bacteriochlorophylls present in the 3Hβ RC (Figure 1(b)). TA data for near-IR probe wavelengths between 850 and 1100 nm are shown in Figure 2(a). This probe wavelength region is particularly important to track the initial steps of charge separation in RCs which, if active, will generate P^+^(B_A_β_A_)^−^; as the product of primary electron transfer. Absorbance changes at 880 nm report the electronic state of P, and bacteriochlorophyll anions have a spectrally distinct absorption band that peaks at ∼1000 nm.^42,43^ The spectra in Figure 2(a) were dominated by a negative feature at 880 nm corresponding to ground state bleaching of P, and stimulated emission from P*. This feature developed quickly in a biexponential fashion and then decayed monoexponentially. Simultaneous fits to multiple probe wavelengths in this region (Figure 2(b)) returned exponential rises of 140 ± 59 fs (93% amplitude) and 2.12 ± 0.49 ps (7%). As 595 nm light excited all five bacteriochlorophylls rather than just the two that form P, we associate the fastest time constant with (B_A_/B_B_/β_A_)* → P* electronic energy transfer,^44–46^ whereas the secondary slower rise (< 7% amplitude) is attributed to a small amount of Crt* → B_B_* → P* electronic energy transfer. The latter minor component was due to overlap of the ∼50 nm FWHM pump pulse with the red-edge of the broad absorbance band of the single 15 *cis-cis’;* spheroidenone in the RC protein^7,47,48^(as evident from carotenoid spectral signatures in the visible probe window– see SI, Section 5). P* then decayed with a time constant of 162 ± 0.2 ps via a mechanism that dominantly must have involved direct transfer back to the ground state as no discernable transient was seen, within the signal-to-noise ratio, that might be associated with formation of B_A_^−^ or β_A_^−^ products. These anions would be expected as a positive shoulder on the ground state bleach/stimulated emission feature at ∼1000 nm. We note that recovery of the ground state bleach was not entirely complete within the 1.9 ns measuring window (see the kinetic trace at 880 nm in Figure 2(b)), with a small offset remaining. This long-lived bleach is likely a signature of a very small percentage of RCs that were able to undergo charge-separation. This combination of spectral features was consistent with the expected effects of combining the triple H-bond^36–38^and β-mutations,^39,40^very strongly reducing the quantum yield of charge separation and greatly increasing the lifetime of the P* excited state from ∼3 ps to over 150 ps.

**Figure 2.**
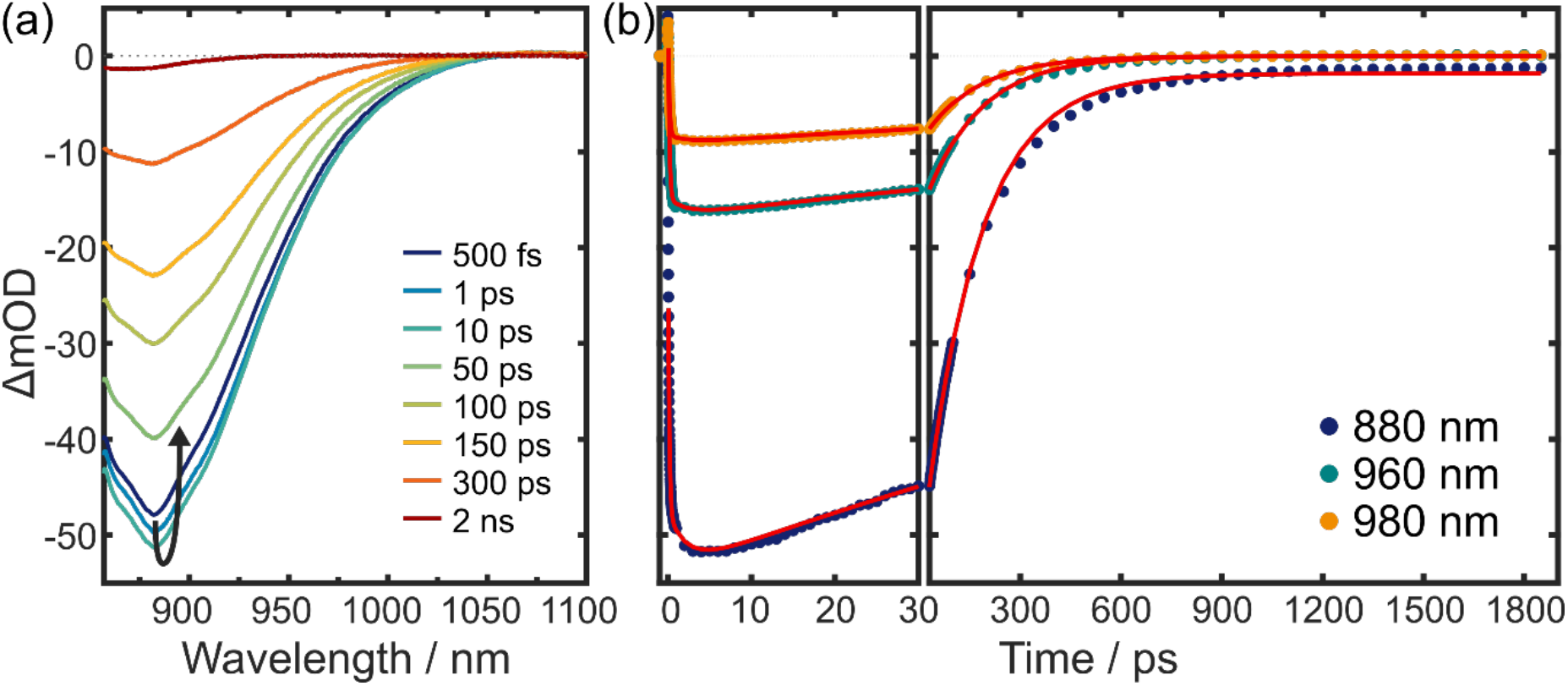
(a) Near-infrared transient absorption spectra of 3Hβ RCs for the displayed pump-probe time delays. (b) Kinetics for three different probe wavelengths and (dots) and fits to the data (solid red lines).

Having characterized the energy and electron transfer dynamics in isolated 3Hβ RCs with TA spectroscopy, we next investigated the excited state dynamics of QD–RC nanoconjugates using TCPSC (see SI, Section 6 for TCSPC experimental details). Fluorescence decay traces for QDs, RCs and four QD–RC biohybrid nanoconjugates with RC:QD molar ratios of between 1:1 and 10:1 are shown in Figure 3. Samples were excited with ∼100 fs 430 nm laser pulses to preferentially excite QDs at a point where background absorbance by RCs was lowest. To achieve different RC:QD ratios the concentration of RCs was fixed and the concentration of QDs varied.

**Figure 3.**
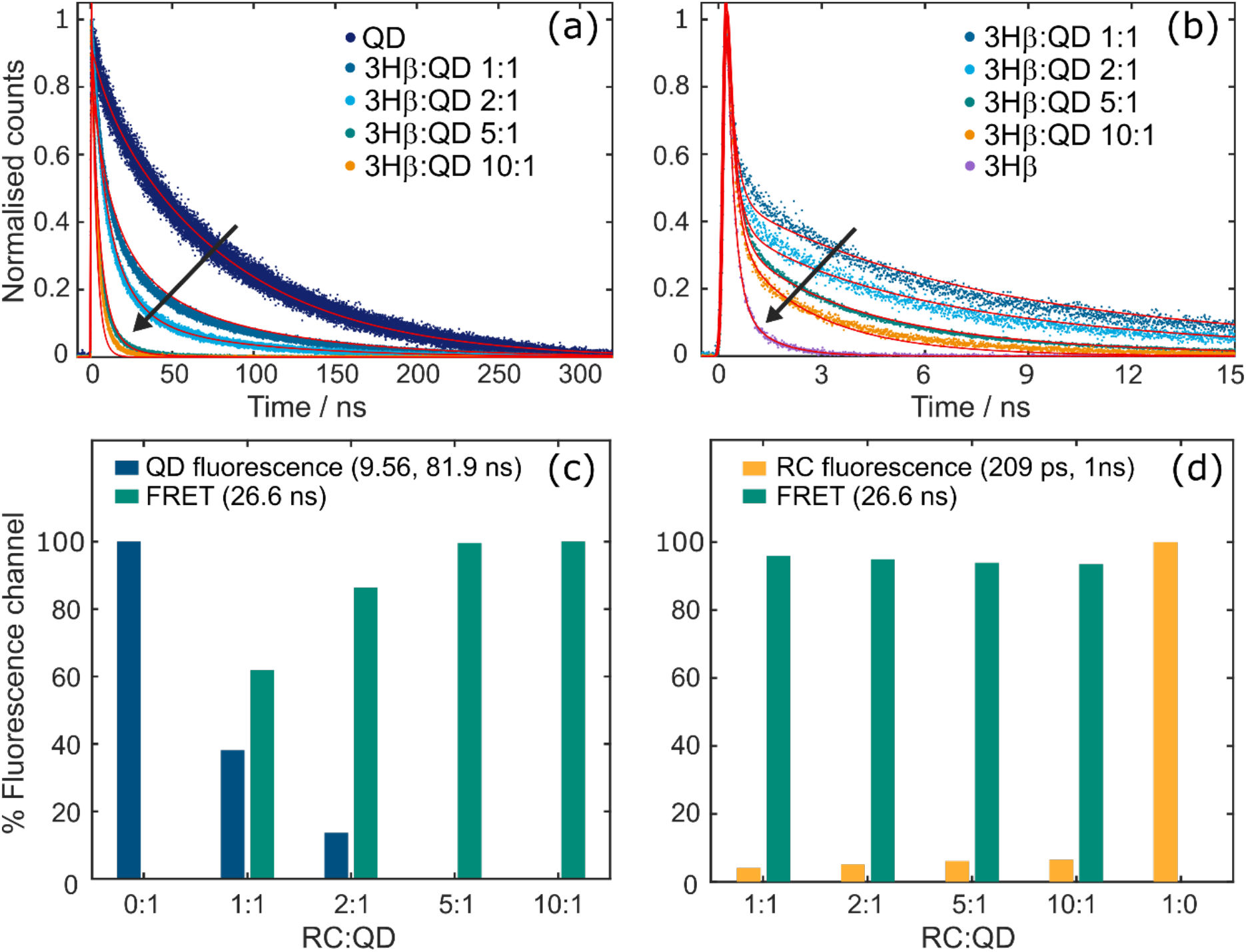
TCSPC analysis of fluorescence decay. (a) Normalized fluorescence data (scatter plots) and overlaid fits returned by global analysis (red solid lines) for QDs and RC–QD conjugates for the peak of QD fluorescence (750 nm). (b) Equivalent data for 3Hβ and RC–QD conjugates at the peak of RC P* fluorescence at 905 nm. In (a) and (b), black arrows indicate the trend upon increasing the RC:QD stoichiometry. (c) Contributions of QD fluorescence (two processes summed) and emission quenched by FRET (two channels summed) to the 750 nm signal for different RC:QD ratios; the 0:1 sample had only QDs. (d) Contributions of intrinsic RC fluorescence (two processes summed) and FRET (two channels summed) to the 905 nm signal at different RC:QD ratios; the 1:0 sample had only RCs. The amplitudes used to determine these percentage contributions are shown in Tables S1 and S2.

Figure 3(a) shows the normalized TCSPC signals collected at 750 nm, which corresponds to the peak of QD fluorescence. Data for the QD-only sample (Figure 3(a), dark blue trace) were fitted with equation S2, comprising a biexponential decay function with a shorter 9.56 ± 0.03 ns component attributed to fluorescence from the excitonic QD state, and a longer 81.89 ± 0.03 ns component. The latter was assigned to delayed fluorescence from the QD excitonic state after de-trapping of localized surface trap states. Approximately 88% of the fluorescence originated from the delayed emission (Figure 3(c)), in line with previous measurements on 6.5 nm CdTe QDs.^49,50^ Time profiles for the QD–RC conjugates (Figure 3(a)) showed a more rapid fluorescence decay that accelerated as the RC:QD stoichiometry increased, consistent with expectations from previous steady-state fluorescence quenching experiments.^31^These data support the hypothesis that FRET occurs between QD and RCs in the conjugate, and that nonradiative FRET within the nanoconjugate competes with the intrinsic QD radiative decay with the lifetimes outlined above. The fluorescence decay traces for the conjugates could be globally fitted with equation S3 (together with data collected at 905 nm, as described below) by inclusion of an additional 26.6 ± 0.1 ns exponential decay component assigned to QD → RC FRET. To minimize the number of free parameters, the QD fluorescence lifetimes and relative amplitudes were both fixed to the above parameters determined from QD-only fluorescence data (see Table S1). The kinetic model also assumed that, regardless of trapping/de-trapping in the QDs, FRET could occur only from the excitonic state of QD with a time constant of 26.6 ns.

The QD:RC stoichiometries shown in Figures 3(a,b) represent average compositions that do not take into account heterogeneity. As demonstrated by Liu *et al.*,^31^each mixture is more appropriately described by a probability distribution of different QD–RC_*n*_ ratios (for *n =* 0–15) as determined by a dissociation constant (*K*_d_) that describes a dynamic equilibrium between RCs bound to QDs and free in solution. This was advantageous for the total FRET efficiency of the nanoconjugates; the more RC acceptors a given QD donor binds, the increased number of possible degenerate EET pathways, which at a macroscopic level boosts the probability of FRET by *n*-fold.^51^The simulated binding probability distributions are given in Figure S3, and were determined by optimizing the value of *K*_d_ in our global kinetic analysis. The *K*_d_ value determined for the 3Hβ RC was 17.3 × 10^−9^M, which was ∼two-fold larger than determined for WT RCs, indicating a slightly weaker binding. Full details of the kinetic model and fitting procedure are given in the SI Section 7.

As summarized in Figure 3(c), even at low RC:QD ratios the relative amplitude of the intrinsic QD fluorescence (blue bars) was outcompeted by the corresponding FRET amplitude (green bars). This is indirect evidence of the effectiveness of the QD RC FRET, as at stoichiometries of 1:1 or 2:1 the subpopulation of QDs with no bound RCs is still significant. Inevitably, increasing the RC:QD ratio increases the probability of FRET, and thus accounts for the observed shortening of the fluorescence decays (orange trace in Figure 3(a)), as slow QD emission was outcompeted by faster FRET. The 26.6 ns FRET pathway dominated the decay in the 5:1 and 10:1 nanoconjugates (Figure 3(c), green bars), peaking at the 5:1 ratio with consequent disappearance of the QD fluorescence channels.

Turning to acceptor fluorescence, Figure 3(b) shows the normalized TCSPC signal collected at 905 nm, a wavelength where RC fluorescence was almost entirely spectrally isolated from QD fluorescence (see Figure 1(b)). For 3Hβ RCs alone (Figure 3(b), violet trace) the data were fitted with equation S4 comprising a biexponential decay function with time constants of 209 ± 4 ps and 1.00 ± 0.04 ns, the nanosecond component comprising 11% of the total fluorescence decay (Table S1). Guided by the near-infrared region TA data, the picosecond component was assigned to fluorescence from P* in RCs that were directly excited by the 430 nm pulse. The minor nanosecond component is assigned to delayed P* fluorescence from a small percentage of directly excited RCs that were able to undergo charge separation and slowly reformed P* through reverse electron transfer. The total fluorescence quantum yield of 3Hβ RCs was estimated to be 0.14% (see SI, Section 3 for analysis). To estimate the initial quantum yield of charge separation for 3Hβ RC, we made the assumption that all RCs able to undergo charge-separation recombined to P* (*i.e.* none recombined to ground state). This analysis returned an initial quantum yield for charge separation of 0.02%.

For the conjugates, as the RC:QD ratio increased the fluorescence profile at 905 nm (Figure 3(b)) displayed the opposite trend to that observed for QD fluorescence at 750 nm (Figure 3(a)). The normalized fluorescence trace for the 1:1 ratio had the slowest decay, and as the RC:QD ratio was increased the rate of decay also increased. The 430 nm pulse did not exclusively excite QDs due to an appreciable absorption cross-section for the 3Hβ RC at this wavelength. As a result, the observed 905 nm fluorescence was due to both direct excitation of RCs and FRET from associated QDs. The relative contributions of these two processes were dictated by the mixture of RC and QD in each sample, with the dynamics tending to resemble those of the RCs alone most strongly in the sample with the highest RC:QD ratio, where the relative absorbance of the RC at 430 nm was highest.

The TCSPC data for the conjugates were globally fit with equation S5. Amplitudes returned from the fits are shown in Tables S1 and S2. All data for RC–QD conjugates required a component to account for FRET from the QDs. Even at the lowest 1:1 stoichiometry most of the fluorescence decay was caused by RCs being excited through FRET (Figure 3(d)). This percentage only slowly declined as the relative amount of RCs increased, due to an increased probability of direct excitation and subsequent fluorescence from the RCs.

The average QD–RC donor-acceptor distance was determined directly using the FRET rate obtained from TCSPC experiments, producing a value of 6.28 ± 0.04 nm (see SI section 8 for details of our analysis). This was close to values of 6.4 – 6.9 nm estimated in our previous study for a mutant RC with just the β-mutation (and therefore the same pigment composition as the 3Hβ RC).^31^The new value is more reliable, as the use of TCSPC enabled direct determination of the FRET rate constant from a global analysis of the fluorescence lifetime data, as opposed to a more complex analysis which required extrapolation. From the Förster rate constant, and knowledge of the QD radiative rates, we also directly derived a FRET efficiency of 0.75 ± 0.01 for a single donor–acceptor pair. This was again close to a value of ∼0.71 estimated in our previous work for 5:1 and 10:1 conjugates formed from RCs with the β-mutation.^31^

We note that the total quantum yield of FRET (<14%) was far lower than the FRET efficiency as it is limited by the ∼20% QD fluorescence quantum yield, as determined by a high nonradiative relaxation quantum yield. A higher FRET quantum yield could potentially be achieved with far more fluorescent QDs, but this is not without its own issues. To increase the fluorescence quantum yield of QDs often requires surface functionalization/passivation to suppresses the number of trap states which predominantly decay non-radiatively and thus lower the fluorescence quantum yield. However, whilst this would increase the direct prompt radiative channel of QDs (9.56 ns), it would also deplete the reservoir of QDs which have delayed fluorescence from the excitonic state after de-trapping of surface states (81.89 ns). Kinetic analysis of QD–3Hβ hybrids revealed the latter channel is the main precursor to FRET (τ = 26.6 ns) in the nanoconjugates (see Table S1), due to the more favorable competition of radiative and non-radiative lifetimes.

A schematic of the competing absorption, non-radiative decay and fluorescence pathways in the 3Hβ RC nanoconjugates is shown in Figure 4(a-c). Depending on the precise RC:QD ratio investigated, 430 nm light can excite either the QD or RC components of the nanoconjugate. Direct RC excitation (Figure 4(a)) leads to a biphasic fluorescence decay due to minimal (∼0.04%) charge separation. Excitation of the QDs can lead to direct or delayed fluorescence from the excitonic state (Figure 4(b)) or FRET to conjugated RCs and subsequent red-shifted 905 nm fluorescence from RCs (Figure 4(c)). The FRET donor–acceptor pair-wise efficiency was determined to be 0.75 ± 0.01. The high FRET efficiency allied with the high surface coverage of QDs with multiple RCs, even at 5:1 RC:QD mixtures, meant that the FRET channel dominated the radiative relaxation dynamics.

**Figure 4.**
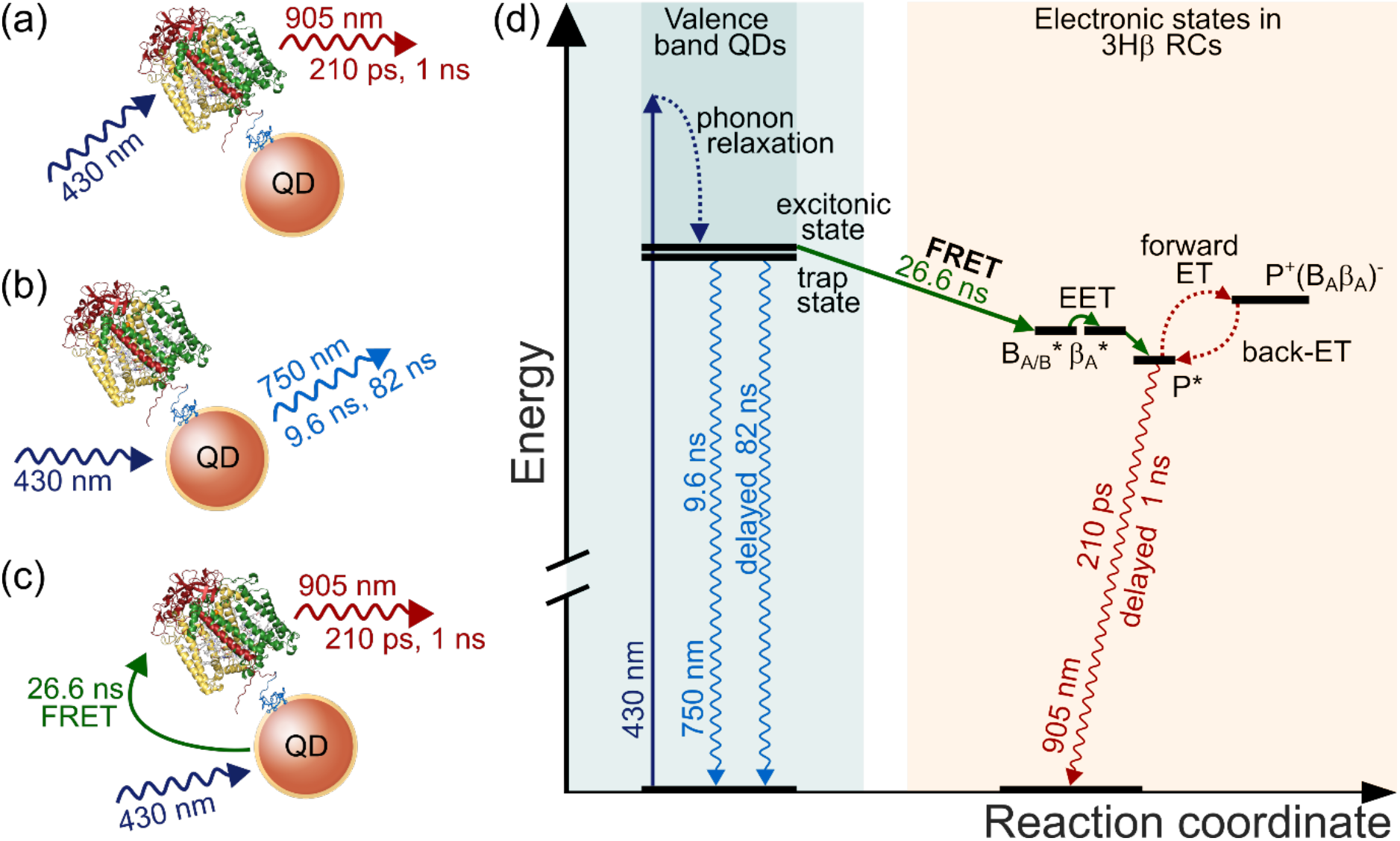
Schematic of radiative and non-radiative decay pathways in 3Hβ RC–QD nanoconjugates, indicating EET and ET components. Measurements on nanoconjugates revealed a mixture of three processes depending on the QD:RC stoichiometry: (a) direct excitation of RCs, and subsequent biexponential fluorescence; (b) excitation of QDs and the associated intrinsic radiative decay from quantum and surface trap states and (c) excitation of QDs, FRET to bound RCs, and fluorescence from those RCs. (d) Energy level diagram with the main radiative and non-radiative pathways associated with QD–3Hβ RC conjugates and their associated time constants. Note FRET to the cofactors of the RC B-branch and subsequent energy transfer to P* is omitted from (d) for the sake of clarity.

In summary, to explore the kinetics and efficiency of energy transfer to purple bacterial RCs from a synthetic antenna complex we engineered a RC with a new combination of mutations that almost completely switches off charge separation, creating a fluorescent energy transfer acceptor. These 3Hβ RCs have a dominant excited state lifetime of 162 ± 0.2 ps, with a fluorescence maximum at 910 nm consistent with emission from the excited state of the primary electron donor. Conjugation of 3Hβ RCs to 6.5 nm diameter CdTe QDs generated an efficient FRET pair, the energy transfer dynamics of which could be directly monitored using TCSPC. FRET occurred within the nanoconjugate with a 26.6 ± 0.1 ns lifetime and a pair-wise efficiency of 0.75 ± 0.01. These parameters enabled a direct determination for the donor-acceptor distance of 6.28 ± 0.04 nm. This combination of natural and synthetic components in a nanoconjugate allows light harvesting to occur with an appreciable quantum yield from the entire visible and near-infrared spectrum, and we envisage the incorporation of additional light harvesting moieties could produce further enhancements that can be characterized using this approach combining ultrafast spectroscopy with the tailoring of photoprotein properties.

## Associated content

Supporting Information is available, providing details of the following: 3Hβ RC mutations, expression, purification and nanoconjugate preparation; steady-state spectroscopic characterization of 3Hβ RCs; TA experimental setup; TA visible probe data of 3Hβ RCs; TCSPC experimental setup; global fitting of TCSPC data sets; FRET quantum yield and distance calculations.

## Notes

The authors declare no competing financial interest.

## Acknowledgements

T.A.A.O acknowledges financial support from the Royal Society for a Royal Society University Research Fellowship (UF1402310, URF\R\201007) and a Research Fellows Enhancement Award (RGF\EA\180076), M.R.J and J.L. acknowledge funding from the EPSRC/BBSRC Synthetic Biology Centre for Doctoral Training (EP/L016494/1) and from the BrisSynBio Synthetic Biology Research Centre at the University of Bristol (BB/L01386X/1). G.A. acknowledge EPSRC for DTP Ph.D. studentship (EP/N509619/1).

---

**TOC Graphic.**
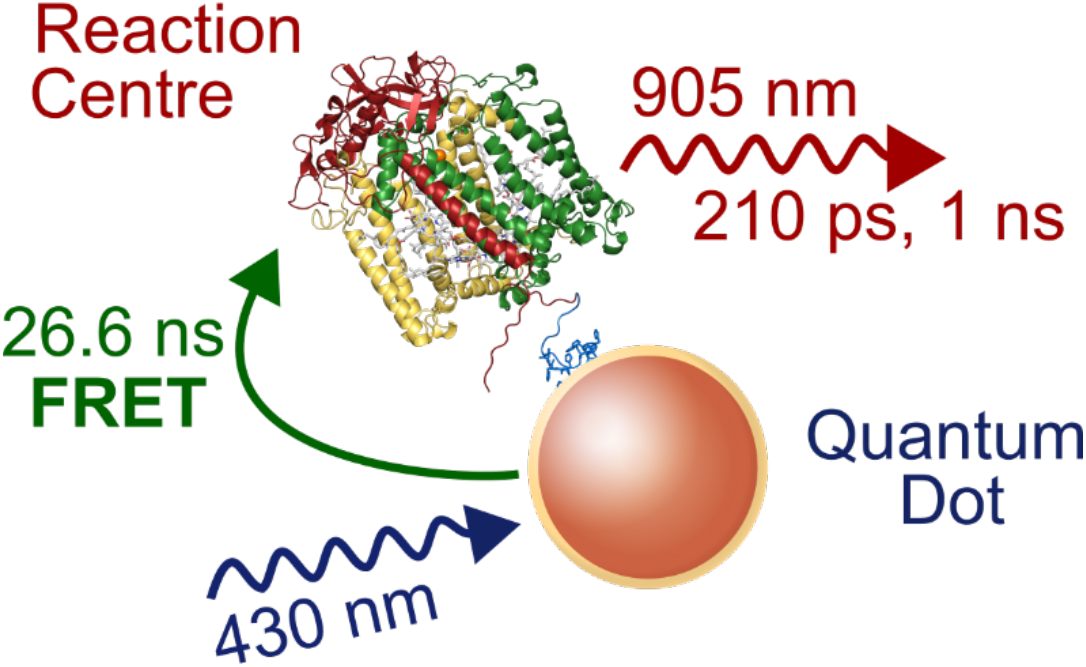

