## Supporting information for "High efficiency excitation energy transfer in biohybrid quantum dot–bacterial reaction center nanoconjugates"

### 1. Engineering of the 3Hβ RC

The 3Hβ RC has four single point mutations. These are: (i) substitution of leucine at position 131 of the L-polypeptide with histidine (LL131H), (ii) substitution of leucine at the position 160 of the M-polypeptide with histidine (LM160H), (iii) substitution of phenylalanine at position 197 of the M-polypeptide with histidine (FM197H), and substitution of leucine at position 214 of the M-polypeptide with histidine (LM214H). A RC with changes (i-iii) has been reported previously; each of the three histidines donates a hydrogen bond to a carbonyl group of one of the two bacteriochlorophylls that make up the primary electron donor P.[1–3] The fourth mutation causes replacement of the second electron acceptor bacteriopheophytin by a bacteriochlorophyll, and is often referred to as the β-mutation.[4,5] Using the QuickChange procedure (Agilent) the four mutations were introduced sequentially into plasmid pv109, which is a derivative of pUC19 containing a 1901 bp XbaI–BamHI restriction fragment encompassing the RC genes *pufLM* modified to place the protein sequence LALVPRGSSAHHHHHHHHHH at the C-terminus of the M-polypeptide.[6] The modified XbaI–BamHI restriction fragment was then shuttled into plasmid pvLMt,[6] and the resulting derivative expressed in *Rba. sphaeroides* strain DD13.[7] Bacterial growth, cell harvesting and purification of the $His_{10}$-tagged 3Hβ RCs was as described in detail in previous studies,[6] as was the assembly of nanoconjugates with different RC:QD molar ratios.[8,9] The latter involved mixing of RCs with water-soluble CdTe QDs that were purchased from PlasmaChem GmbH. These had an emission maximum at 750 ± 5 nm and were coated with 3-mercaptopropionic acid.

### 2. Steady-state spectroscopy of 3Hβ RCs

Figure S1 shows an overlay of absorption spectra of the 3Hβ and WT RCs. Bands in the spectrum of the 3Hβ RC can be assigned by reference to those in the spectrum of WT RCs and known effects of the four mutations. For the WT RC, the bands peaking at 360 and 390 nm are assigned to the Soret bands of the four bacteriochlorophylls and two bacteriopheophytins. The single 15 *cis-cis*' spheroidenone carotenoid contributes to a broad absorption between ~450 and ~600 nm. The sharper features in this region correspond to the $Q_y$ bands associated with two bacteriopheophytins (between 520–550 nm) and four bacteriochlorophylls (at 600 nm). The spectrum of the 3Hβ RC showed the loss of a band at ~545 nm and the appearance of additional absorbance around 600 nm, consistent with the replacement of a bacteriopheophytin by a bacteriochlorophyll. In the WT RC the major bands in the near-IR region are attributable to the two bacteriopheophytins ($H_A$/$H_B$, at 760 nm), the two monomeric bacteriochlorophylls ($B_A$/$B_B$, at 802 nm) and the two strongly excitonically-coupled primary electron donor bacteriochlorophylls (P, at 870 nm). As expected, the band associated with bacteriopheophytin at 760 nm was lowered in intensity in the spectrum of the 3Hβ RC, with the appearance of additional absorption on the blue side of the 802 nm band. Again, this was consistent with the replacement of a bacteriopheophytin by a bacteriochlorophyll.[4,6]

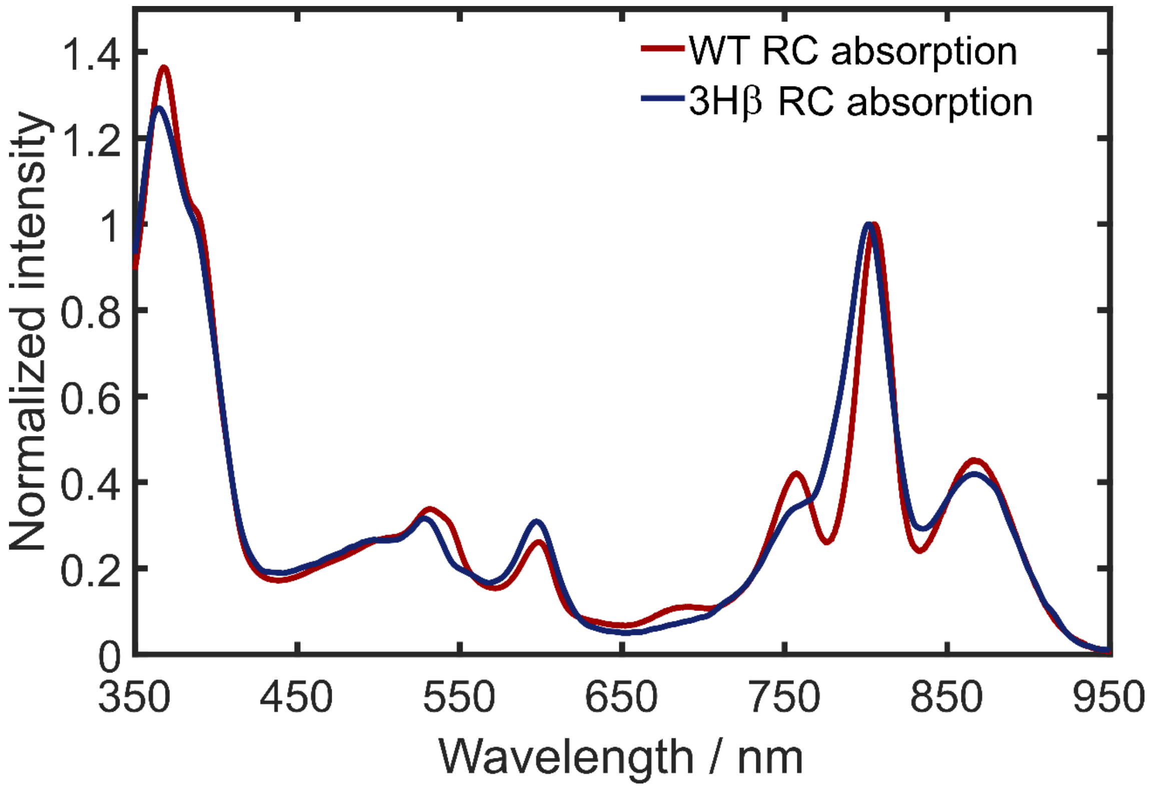

**Figure S1.** Absorption spectra of wild-type RCs (red) and 3Hβ RCs (dark blue), normalized to the ~800 nm band.

### 3. 3Hβ RC fluorescence quantum yield

To determine the fluorescence quantum yield of 3Hβ RCs, the fluorescence of the near-infrared dye IR140 in dimethyl-sulfoxide, which has a well characterized fluorescence quantum yield, was recorded under the same excitation conditions as used to acquire the fluorescence of 3Hβ RCs. Using a previously described proceedure,[10] the fluorescence quantum yield, $\Phi_f$, of the 3Hβ RCs was determined using IR140 as a calibrant via the equation:

$$\Phi_f(3H\beta) = \Phi_f(IR140)\frac{{n_{3H\beta}}^2}{{n_{IR140}}^2}\frac{F_{3H\beta}}{F_{IR140}}\frac{1-10^{-A(595)_{IR140}}}{1-10^{-A(595)_{3H\beta}}} \qquad \text{(S1)}$$

where $F$ is the integrated fluorescence intensity, $A(\lambda)$ is the absorbance at excitation wavelength $\lambda$, and $n$ is the refractive index of the solvent. Using the known fluorescence quantum yield of 0.20 for IR140 in dimethyl-sulfoxide,[11] the fluorescence quantum yield for the 3Hβ RC was estimated to be 0.0014.

### 4. Transient Absorption Spectroscopy

Visible and near-infrared TA experiments were performed using an apparatus built in our laboratory. Part (40%) of the 800 nm fundamental laser output of a 1 W, 1 kHz, Ti:Sapphire amplifier (Coherent, Libra) was divided with a 90:10 beam splitter to generate pump and probe pulses, respectively. The 595 nm pump pulses with a FWHM of ~50 nm were generated using a non-collinear optical parametric amplifier (NOPA).[12] Second order group velocity delay was compensated for by chirped mirrors (Layertech 148545) and fused silica wedges (FemtoOptics, Newport). The pulse duration was determined by a polarization gated frequency-resolved optical gating setup to be 59 ± 5 fs. The pump pulses were chopped at 500 Hz and delayed with respect to the probe via a motorized high precision delay stage (Physik Instrumente, M-531.DG1) with maximum delay of 1.9 ns. The pump pulses were focused into the sample to a spot-size of 100 μm diameter using a plano-concave mirror ($f$ = 20 cm).

Probe pulses were generated by sending a portion of the fundamental laser into either a Sapphire crystal for a visible probe (450 nm to 700 nm) or a YAG crystal for a near-infrared probe (860 nm to 1200 nm). The probe beam was then focused into the sample, recollimated with the collinear signal, and for visible TA measurements spectrally filtered to remove residual fundamental 800 nm light using a short-pass dichroic mirror (DMSP750B, Thorlabs). Finally, the probe and signal were focused into a spectrograph (Shamrock 163, Andor) and detected with a linear 1024 element CCD array detector (Entwicklungsbüro Stresing).

Prior to data collection, the pump fluence was attenuated to ∼16 nJ at the sample to obtain a power density of 2.54 GW $cm^{-2}$, in line with previous RC TA studies.[13] TA measurements were performed under the magic angle condition. The 3Hβ RC samples were freshly prepared and purified prior to TA measurements and diluted to obtain an absorption of ~0.34 at 600 nm. The sample was flowed continuously throughout measurements in a flow cell with a 1 mm path length (Starna, Type 48). Each data cycle was acquired using a randomized sequence of pump-probe time delays.

### 5. Visible region TA of the 3Hβ RC

TA difference spectra for the 3Hβ RC using 600 nm excitation and a visible white light probe supercontinuum (450–720 nm) are displayed in Figure S2. Unlike prior measurements of R26 RCs (a *Rba. sphaeroides* RC that lacks the carotenoid cofactor) there were no obvious features associated with bacteriochlorin anions, which in this region are expected to be present at 630 and 640 nm,[13,14] and in accord with our near-IR probe data shown in the main manuscript (Figure 2). These results further demonstrate that the charge-transfer is greatly disfavored in the 3Hβ RC. The broad bleach between 475 and 565 nm is associated with the 15 *cis*-*cis*' spheroidenone, as the 595 nm (FWHM = 50 nm) excitation catches the long-wavelength tail of its absorption. The visible region TA spectrum is far more complex to decompose compared to the equivalent near-IR spectrum, due to overlapping excited state absorption bands and multiple ground state bleaches from the $Q_x$ bacteriochlorin bands.

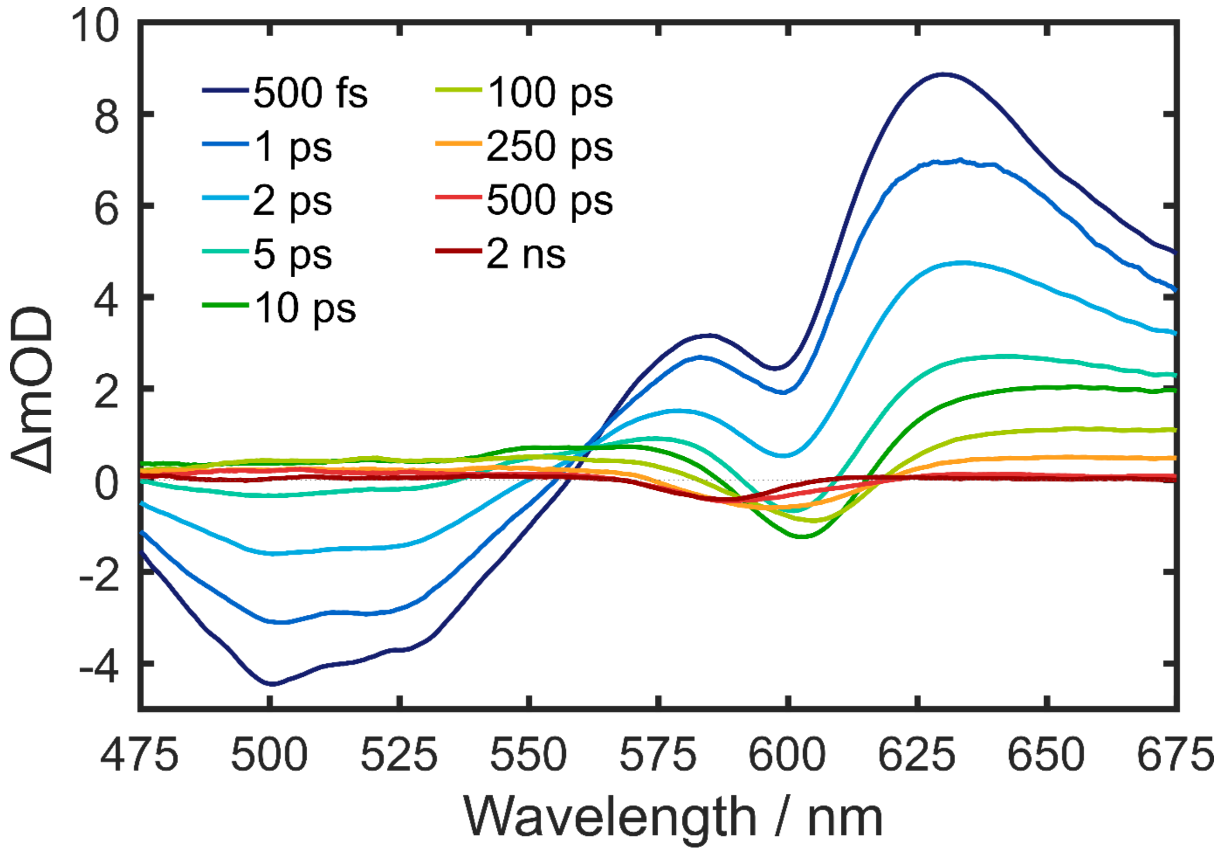

**Figure S2.** Transient absorption spectra of the 3Hβ RC for visible probe wavelengths.

### 6. Time-correlated single photon counting (TCSPC)

Time-correlated single photon counting (TCSPC) measurements were recorded using a homebuilt spectrometer. The 860 nm fundamental laser output of a narrowband high power Ti:Sapphire oscillator (3.7 W, 80 MHz, Chameleon Ultra II, Coherent) was focused in a 2 mm thick BBO crystal ($\theta$ = 29.2°, Eksma), and frequency doubled to 430 nm. Residual near-IR fundamental light was removed using two dichroic beam splitters (Separator 106160, Layertec). To avoid re-excitation of samples, the repetition rate of the laser pulse train was reduced to 3 MHz using a pulse picker (APE cavity dumper kit). The diffracted light from the acousto-optic modulator (AOM) was subsequently focused through a 10 μm pinhole to remove any of the un-diffracted parent beam. The laser light (10 μW) was subsequently collimated and focused into the sample with a second lens. Fluorescence was collected from a 1 cm pathlength sample cuvette at 90º relative to the excitation laser with an infinity corrected microscope objective (4×/0.2 NA Plan Apochromat, Nikon). The fluorescence was filtered to remove any laser scatter and collected at 750 nm (using a 10 nm FWHM bandpass filter; 87-889, Edmund Optics) or at 905 nm (using a 25 nm FWHM bandpass filter; FL905-25, Thorlabs). A polarizer before the detector was set to magic angle relative to the excitation laser to eliminate rotational anisotropy effects. The filtered fluorescence was focused onto an avalanche-photodiode detector (ID100-20-REG, IDQ) with an achromatic doublet (AC508-075-A, Thorlabs). The photon counts from the detector were acquired by a time-to-digital converter (Time Tagger 20, Swabian Instruments) and binned into histograms of 10 ps bins. TCSPC data were acquired using customized LabVIEW software (National Instruments). The instrument response function for the TCSPC experiment was determined to be 170 ps using a colloidal silica solution. TCSPC traces were fitted in MATLAB (MathWorks) to analytical solutions of a gaussian instrument response convoluted with two or more exponential functions. All measurements were performed at room temperature (20 °C).

### 7. Global fitting and modelling of TCSPC dataset

The TCSPC traces for QDs, RCs and nanoconjugates were fitted to the analytical solution of a gaussian instrument response function convolved with multiple exponential decays. For isolated QDs, the fluorescence signal at 750 nm, was modelled via:

$$I_{QDs(750)} = A_1 e^{-k_{D1}t} + A_2 e^{-k_{D2}t}, \quad \text{(S2)}$$

where $k_{D1}$ and $k_{D2}$ are the rate constants associated with the direct and delayed fluorescence from the QD excitonic state, respectively, and $A_1$ and $A_2$ are the associated amplitudes. For the sake of simplicity, equation S2 and those that follow exclude the convolution function.

To account for the heterogeneity in the RC:QD stoichiometry for different donor–acceptor mixtures, we used the deterministic binding model defined by Liu *et al.,*[8] which resulted in a similar macroscopic dissociation constant of 17.3 $\times 10^{-9}$ M, see below. This model accurately predicted the RC:QD stoichiometry probability distribution for WT RCs and QDs in our previous study. As the 3Hβ protein still binds through to QDs via His-tags and in very similar structural way to WT RCs, we expected the same model to accurately predict the 3Hβ–QD binding distributions. The binding distributions for the ratios studied are given in Figure S3, and the associated probabilities ($p(i)$) were used in kinetic modelling of nanoconjugate fluorescence lifetime data.

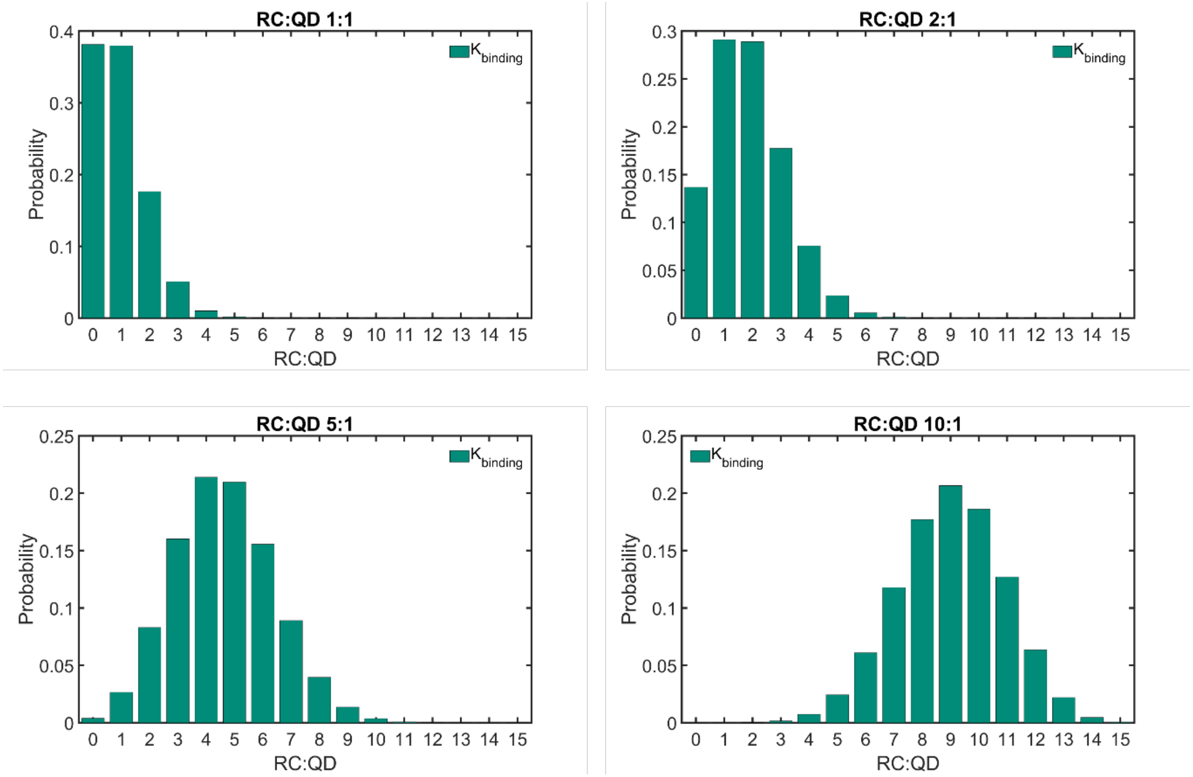

**Figure S3.** Calculated probability distributions for the RC:QD stiochiometry based on the following RC:QD mixture ratios: (a) 1:1, (b) 2:1, (c) 5:1 and (d) 10:1.

As is evident from Figure S3(a), even at the lowest concentration mixtures the probability that a single QD binds multiple RCs is > 20%, meaning there will multiple degenerate QD→RC FRET pathways in each sample.

Based on the calculated stoichiometry probability distributions for the mixtures studied, it was apparent that the fluorescence decay profiles needed to account for a sum between 0–15 RCs (truncated to 15 because the probability of binding 15 RCs in the mixtures studied is negligible), where the pre-exponential terms, ($p(i)$) were determined by the probability distribution above. To account for the multiple possible FRET pathways, a multiplicative factor (equal to specific stoichiometry in the sum) for the FRET rate constant, $k_{FRET}$ was included for both the direct ($k_1$) and delayed ($k_2$) QD fluorescence donor states. Accounting also for some fraction of QDs without RCs tethered, this yielded:

$$I_{conj(750)} = A_3(\sum_{i=0}^{15} p(i)\, e^{-(k_{D1}+ik_{FRET})t}) + \\ + \frac{A_2}{A_1} A_3 \sum_{i=0}^{15} p(i)\, e^{-(k_{D2}+ik_{FRET})t}), \qquad \text{(S3)}$$

Where the ratio between the fluorescence originating QDs (direct *vs.* delayed) was constrained by the ratio determined in QD-only data and $A_3$ is the signal amplitude in nanoconjugate traces.

The isolated signal from 3Hβ RCs at 905 nm was described by a biexponential decay:

$$I_{RCs(905)} = A_4 e^{-k_{A1}t} + A_5 e^{-k_{A2}t}, \qquad \text{(S4)}$$

where $k_{A1}$ and $k_{A2}$ were the rate constants associated with direct fluorescence from P* and delayed fluorescence from back electron transfer and reformation of P*, and associated pre-exponential factors $A_4$ and $A_5$.

The time-resolved fluorescence at 905 nm for the nanoconjugates were also modelled to include the heterogeneity of the binding, the presence of two FRET channels, and the emission from two different RCs pathways:

$$\begin{aligned} I_{conj(905)} = {} & A_4 e^{-k_{A1}t} + A_5 e^{-k_{A2}t} + \\ & + A_6 \sum_{i=1}^{15} p(i) \frac{k_{FRET}}{k_{D1}+ik_{FRET}-k_{A1}} \left(e^{-k_{A1}t} - e^{-(k_{D1}+ik_{FRET})t}\right) + \\ & + A_7 \sum_{i=1}^{15} p(i) \frac{k_{FRET}}{k_{D2}+ik_{FRET}-k_{A1}} \left(e^{-k_{A1}t} - e^{-(k_{D2}+ik_{FRET})t}\right) + \\ & + \frac{A_5}{A_4} A_6 \sum_{i=1}^{15} p(i) \frac{k_{FRET}}{k_{D1}+ik_{FRET}-k_{A2}} \left(e^{-k_{A2}t} - e^{-(k_{D1}+ik_{FRET})t}\right) + \\ & + \frac{A_5}{A_4} A_7 \sum_{i=1}^{15} p(i) \frac{k_{FRET}}{k_{D2}+ik_{FRET}-k_{A2}} \left(e^{-k_{A2}t} - e^{-(k_{D2}+ik_{FRET})t}\right), \qquad \text{(S5)} \end{aligned}$$

Equations S4 and S5 were used to globally fit the QD–RC data for the two different probe wavelengths (750 nm for QDs and 905 nm for RCs). These fits allowed $k_{FRET}$ to float, as well as the amplitude terms $A_{3,6,7}$, and $K_D$ which determined the binding probability distribution and hence determined the $p(i)$ pre-factors. The time constants and the amplitudes associated with radiative decay from RCs or QDs were fixed based on the analysis of the isolated components and fitting to equations S2 and S5, respectively. The fit returned a value of $\tau_{FRET}$ = 26.6 ± 0.1 ns. The amplitudes returned from this analysis are given in Table S1 and Table S2, respectively for the signal at 750 and at 905 nm. Fits to data are shown in the main manuscript.

**Table S1.** Amplitudes associated with the globally fit time constants for different samples at 750 nm and used to obtain the bar plot in Figure 3(c).

| QDs fluorescence (750 nm) | | | | |
|---|---|---|---|---|
| Relative amplitudes (%) | | | | |
| RC:QD Ratio | $A_1p(0)$ (QDs direct $\tau$ = 9.56 ns) | $A_2p(0)$ (QDs delayed $\tau$ = 81.89 ns) | $A_3(1-p(0))$ (QDs direct + FRET $\tau$ = 9.56 + 26.6 ns) | $(A_2/A_1)A_3(1-p(0))$ (QD delayed + FRET $\tau$ = 81.89 + 26.6 ns) |
| 0:1 | 12.06 | 87.94 | — | — |
| 1:1 | 4.60 | 33.55 | 7.46 | 54.39 |
| 2:1 | 1.65 | 12.03 | 10.41 | 75.91 |
| 5:1 | 0.05 | 0.35 | 12.01 | 87.59 |
| 10:1 | 0.00 | 0.00 | 12.06 | 87.94 |

**Table S2.** Amplitudes associated with the globally fit time constants for different samples at 905 nm and used to obtain the bar plot in Figure 3(d).

| RCs fluorescence (905 nm) | | | | | | |
|---|---|---|---|---|---|---|
| RC:QD Ratio | $A_4$ (RCs direct $\tau$ = 209 ps) | $A_5$ (RCs delayed $\tau$ = 0.99 ns) | $A_6$ (FRET to P* from QDs direct) | $A_7$ (FRET to P* from QDs delayed | $(A_5/A_4)A_6$ (FRET to $P^+(B_A\beta_A)^-$ from QDs direct) | $(A_5/A_4)A_7$ (FRET to $P^+(B_A\beta_A)^-$ from QDs delayed) |
| Relative amplitudes (%) | | | | | | |
| 1:0 | 87.21 | 12.79 | — | — | — | — |
| 1:1 | 3.54 | 0.52 | 53.08 | 30.58 | 7.78 | 4.49 |
| 2:1 | 4.45 | 0.65 | 52.51 | 30.26 | 7.70 | 4.44 |
| 5:1 | 5.33 | 0.78 | 51.95 | 29.93 | 7.62 | 4.39 |
| 10:1 | 5.72 | 0.84 | 51.70 | 29.79 | 7.58 | 4.37 |

### 8. FRET efficiency and distance

To determine the FRET pair efficiency, the weighted average lifetime of the donor was first calculated using:

$$\tau_D = \frac{\sum \alpha_i \tau_i^2}{\sum \alpha_i \tau_i},$$

which returned an associated radiative rate constant of 1.24 $\times 10^7$ $s^{-1}$. Together with the FRET rate constant, the efficiency was calculated via:

$$E_{FRET} = \frac{k_{FRET}}{k_{FRET}+k_D} \,. \qquad \text{(S6)}$$

This analysis returned a value of $E_{FRET}$ of 0.75 ± 0.01.

The FRET rate constant also allowed direct calculation of the Förster distance:[15]

$$R^6 = 8.785 \times 10^{-25} \frac{\Phi_f(D)}{\tau_D} \frac{1}{k_{FRET}} \frac{\kappa^2 I}{n^4}\,, \qquad \text{(S7)}$$

where $\Phi_f(D)$ is the fluorescence quantum yield of the isolated QD donor (0.197), $\kappa$ is the orientation factor between the dipoles (here $\kappa^2$ is assumed to be 2/3), $I$ is the Förster spectral overlap integral of QD fluorescence and 3Hβ RCs absorption (4.61 $\times 10^{-12}$ $M^{-1}$ $cm^3$), $\tau_D$ is the fluorescence lifetime of the isolated QDs, and $n$ is the refractive index of the medium (this was chosen to be that of water buffer solution, 1.33). The Förster spectral overlap integral was obtained using software freely available at FluorTools.com. This calculation returned a FRET distance of 6.28 ± 0.04 nm.
